# Drastic differential CRISPR-Cas9 induced mutagenesis influenced by DNA methylation and chromatin features

**DOI:** 10.1101/2022.02.28.482333

**Authors:** Trevor Weiss, Peter A. Crisp, Krishan M. Rai, Meredith Song, Nathan M. Springer, Feng Zhang

## Abstract

CRISPR-Cas9 mediated genome editing has been widely adopted for basic and applied biological research in eukaryotic systems. While many studies considered DNA sequences of CRISPR target sites as the primary determinant for CRISPR mutagenesis efficiency and mutation profiles, increasing evidence revealed the significant role of chromatin context. Nonetheless, most of the prior studies were limited by the lack of sufficient epigenetic resources and/or by only transiently expressing CRISPR-Cas9 in a short time window. In this study, we leveraged the wealth of high-resolution epigenomic resources in *Arabidopsis thaliana* to address the impact of chromatin features on CRISPR-Cas9 mutagenesis using stable transgenic plants. Our results indicated that DNA methylation and chromatin features could lead to significant variations in mutagenesis efficiency by up to 250 folds. Low mutagenesis efficiencies were mostly associated with repressive heterochromatic features. This repressive effect appeared to persist through cell divisions but could be alleviated through substantial reduction of DNA methylation at CRISPR target sites. Moreover, specific chromatin features, such as H3K4me1, H3.3, and H3.1, appear to be associated with significant variations in CRISPR-Cas9 mutation profiles reflected by the 1 bp insertion rates. Our findings provided strong evidence that specific chromatin features could have significant and lasting impacts on both CRISPR-Cas9 mutagenesis efficiency and DNA double strand break repair outcomes.

## Introduction

CRISPR-Cas based genome editing technologies have greatly advanced both basic and applied biological research. Among them, CRISPR-Cas9, the class II bacterial CRISPR system, has been the first and the most widely adopted in eukaryotes (Jinek et al. 2012). The key steps in CRISPR-Cas9 mediated genome editing involve searching, binding and then cleaving a 20-nucleotide target site directed by a guide RNA (gRNA). The resulting cleavage product with double-strand breaks (DSBs) can then be repaired by either error-prone DNA repair pathways, such as classical non-homologous end-joining (NHEJ) or microhomology-mediated end-joining (MMEJ), or by a template DNA-dependent pathway, *i*.*e*. homology directed repair (HDR) (Chen et al. 2019). Thus, specific mutations, including insertions, deletions or point mutations, can be introduced by employing distinct DNA repair machineries (Chen et al. 2019). Previous studies indicated that the CRISPR targeted sequence is the primary determinant for mutagenesis efficiency and mutation profile (Allen et al. 2018; Lazzarotto et al. 2020). Several tools have been developed to predict the efficiency and mutation outcomes solely based on CRISPR targeted sequences (Allen et al. 2018; Concordet and Haeussler 2018; Xiang et al. 2021). However, the predictability of these tools, primarily based on the large dataset from human cells, often varies and appears to translate poorly to other species, such as plants (Naim et al. 2020). This observation suggested that non-sequence features could influence CRISPR-Cas9 mutagenesis.

Increasing evidence revealed negative correlations between CRISPR-Cas9 mutagenesis rates and heterochromatic signatures or low chromatin accessibility in multiple systems, such as yeast, zebrafish, mouse, human and rice (Daer et al. 2017; G. Liu et al. 2019; Uusi-Mäkelä et al. 2018; Wu et al. 2014; Yarrington et al. 2018). However, most of these studies were conducted at various genomic locations, making it difficult to separate the effect of chromatin context from those of DNA sequences. Recently two studies investigated the impact of chromatin features by using over 1,000 copies of integrated reporter sequences to fix the sequence variables (Gisler et al. 2019; Schep et al. 2021). Their findings confirmed the previous observations that heterochromatin has a negative impact on CRISPR-Cas9 mutagenesis efficacy. Notably, specific chromatin features were also identified to impact both efficiency and mutation outcomes (Gisler et al. 2019; Schep et al. 2021). Nevertheless, these studies were limited by two factors: 1) the bias of the integration sites in certain genomic locations, and 2) the ambiguity from whether the newly integrated sequences can quickly and faithfully adopt the local chromatin context. Furthermore, most of these previous studies were conducted in cell lines within a short time window (usually less than 72 hours) using transiently expressed CRISPR-Cas9. It is still unclear whether the heterochromatic features have a long-lasting effect on CRISPR-Cas9 mutagenesis, or only delay it (Kallimasioti-Pazi et al. 2018).

In this study, we leveraged the high resolution epigenomic resources in the model plant species, *Arabidopsis thaliana*, to address the impact of chromatin features on both CRISPR-Cas9 mutagenesis efficiency and mutation outcomes. To fix the sequence variable, the Arabidopsis genome was scanned to identify identical CRISPR target sites located in various chromatin contexts. By using stable CRISPR-Cas9 transgenic plants targeting multiple chromosomal regions, we systematically characterized mutagenesis efficiency and mutation outcomes with 23 distinct DNA methylation and chromatin features using a Next Generation Sequencing (NGS) approach. Our findings provided insight into the influences of chromatin features on CRISPR-Cas9 mutagenesis and DNA repair outcomes. This could help develop better technologies for more efficient and precise genome editing.

## Results

### Identification of identical CRISPR-Cas9 sites in diverse chromatin contexts

To identify identical CRISPR target sites in various chromatin contexts, the Arabidopsis *Col*-0 reference genome was scanned for 20 bp CRISPR-Cas9 recognition sequences with 3 bp NGG (the PAM sequence) at the 3’ end. Out of 7,376,476 distinct target sites, 19,161 were identified as repeating 7-25 times across the genome (Figure 1A). A series of filters were then applied to remove the sites with one of the following features: simple repeats, GC content outside the range of 40-60%, matching sequences in mitochondria or chloroplast genomes, or containing no overlapping restriction enzyme site for subsequent mutation genotyping. The remaining 7,971 sequences, representing 92,117 total genomic sites, were assessed for three key chromatin features using 100 bp windows: chromatin accessibility indicated by ATAC-seq scores (Lu et al. 2016), DNA methylation patterns categorized as DNA methylation domain (RdDM, heterochromatin, CG-only, unmethylated and intermediate) (Crisp et al. 2017; Springer and Schmitz 2017), and nine chromatin states (Supplemental Dataset 1) (Sequeira-Mendes et al. 2014). Seven multicopy CRISPR sites (MCsites) were identified with individual sequences in each family having highly diverse chromatin contexts, including both open and closed chromatin, at least 3 different DNA methylation domains, and at least 2 distinct chromatin states (Figure 1B-C; Supplemental Dataset 2).

**Figure 1.**
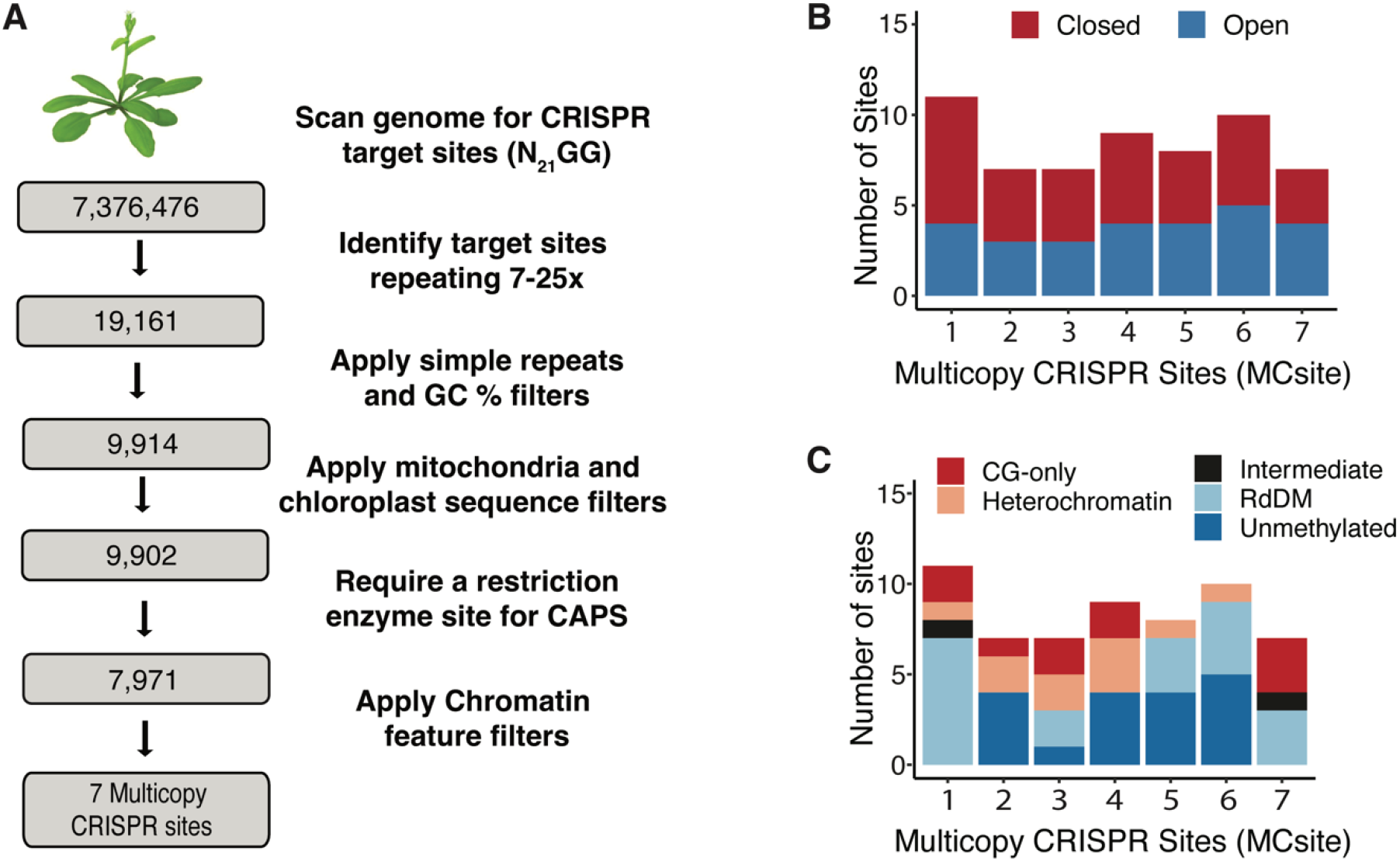
Identification of multicopy CRISPR sites (MCsites) in various chromatin contexts. *(A)* Bioinformatic pipeline used to identify MCsites. *(B)* Characterization of the chromatin accessibility as Closed (red) or Open (blue) at each MCsite using ATAC-Seq data. *(C)* Characterization of DNA methylation domains at each MCsite. CG-only (red) sites contain greater than 40% mCG; Heterochromatin (tan) sites contain greater than 40% mCG and mCHG methylation; RdDM (light blue) sites contain mCG, mCHG, and mCHH, with at least 15% mCHH; Unmethylated (blue) sites contain less than 10% mCG, mCHG, and mCHH; Intermediate (black) sites are everything else with data that didn’t meet any of the above criteria.

### Differential CRISPR-Cas9 efficiencies were associated with distinct chromatin features

Next, we evaluated CRISPR-Cas9 efficacy for each of the seven MCsite families. T-DNA constructs, containing a CRISPR-Cas9 expression cassette, a firefly luciferase reporter and a BAR selection marker gene, were made to target each MCsite (Supplemental Figure S1A). In each construct, the CRISPR single guide RNA (sgRNA) cassette consisted of 2 guide RNAs (gRNAs), one targeting the MCsite and the other targeting a single-copy endogenous gene, Cheletase2 (CHLl2), as the CRISPR mutagenesis control (Figure 2A) (Mao et al. 2013). It is worth noting that we intentionally chose the CaMV 35S promoter to drive expression of the Cas9 and gRNA sequences because this promoter has much lower activity in Arabidopsis embryos than leaves (Wang et al. 2015; Yan et al. 2015). By reducing the mutagenesis potential in early embryo development stages, we were able to capture more independent mutation events in somatic cells during leaf development.

**Figure 2.**
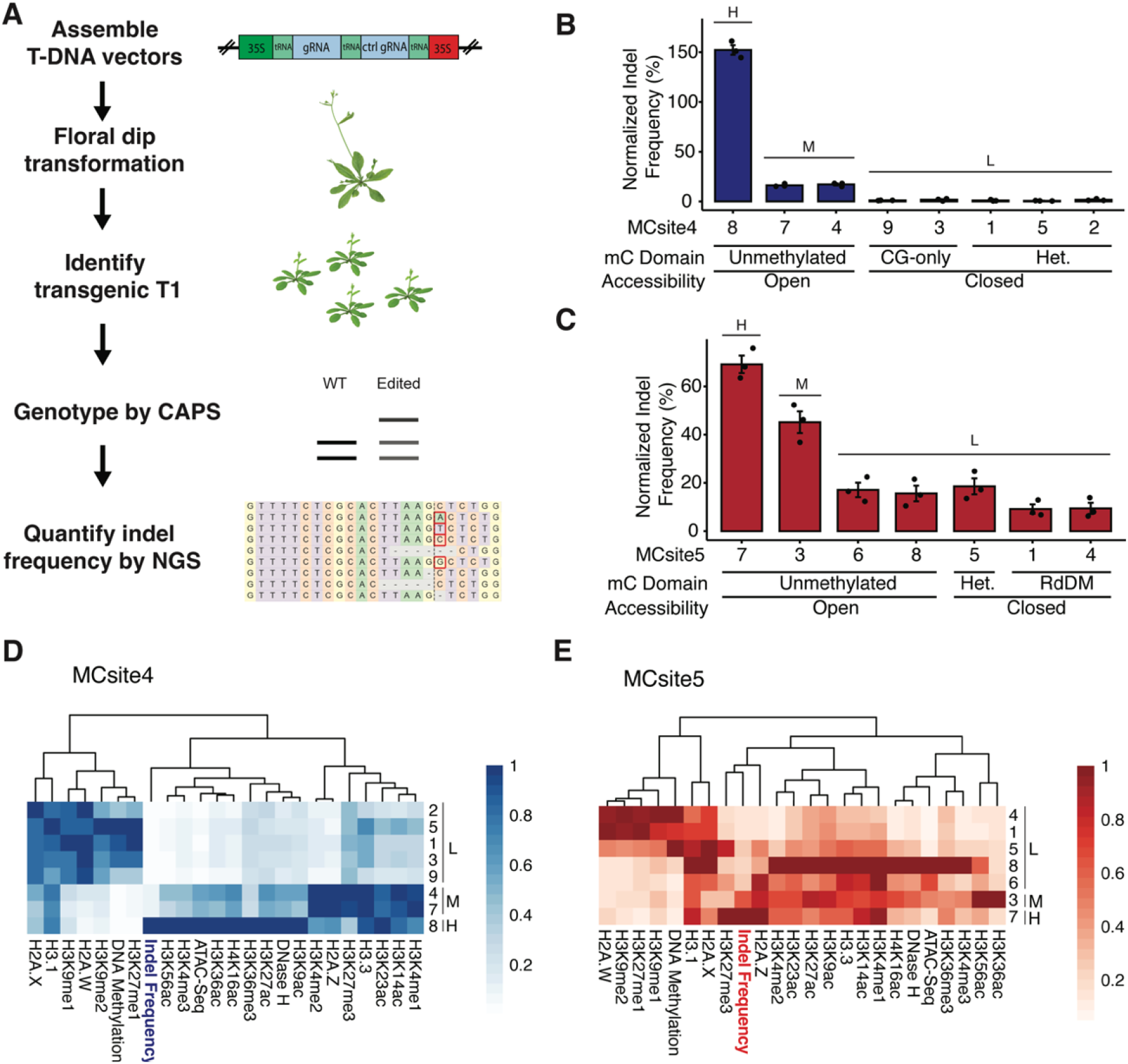
Characterization of CRISPR-Cas9 mutation frequencies and chromatin features at MCsites. *(A)* The NGS-based workflow to analyze CRISPR-Cas9 mutagenesis efficiency and repair profile at MCsites. *(B)* and *(C)* Normalized mutation frequency for individual MCsite4 and MCsite5 CRISPR targets. The standard error (SEM) was calculated for each target site with three replicates. A one-way analysis of variance and Tukey’s multiple comparison test were performed to group the target sites into H (high), M (moderate), and L (low) mutagenesis levels. The DNA methylation domain and accessibility features were listed below each target site. Het stands for Heterochromatin DNA methylation domain. *(D)* and *(E)* Hierarchical clustering heatmap of distinct chromatin features (x-axis) for individual MCsite4 or MCsite5 sites (y-axis). H (high), M (moderate), and L (low) indicated the mutagenesis group for each site. Each feature was normalized to a scale of 0-1, with 1 indicating the highest level (the darkest color) and 0 indicating the lowest level (white). The dendrogram indicated similarity clustering of chromatin features.

After transformation of each T-DNA construct, the resulting CRISPR-Cas9 transgenic plants (T1) were first assessed for mutagenesis efficiency at the CHLl2 control site using the Cleaved Amplified Polymorphic Sequences (CAPS) method. Individual plants with detectable mutagenesis at the control site were then analyzed at each target site. Using the CAPS assay we were able to identify two MCsites, MCsite4 and MCsite5, that produced notable mutagenesis for at least one of the CRISPR target sites (Supplemental Figure S1B). MCsite4 and 5, totaling 15 individual sites, were found spanning all 5 chromosomes (Supplemental Figure S1C). Among the 15 sites, four seemed to overlap with Arabidopsis genes, and only 1 site, MCsite5.8, was detected in transcribed RNA sequences (Supplemental Dataset 2).

CRISPR-Cas9 mutagenesis efficiency was then assessed at the individual sequences of MCsite4 and 5 by quantifying indel mutations using the NGS assay. Consistent with the observation from the CAPS assay, substantial variations were observed across individual target sites (Supplemental Figure S2A). To control for the variation in CRISPR-Cas9 expression levels between plants, the mutation rate at each targeted site was normalized to the CHLI2 control. As a result, the normalized frequencies at MCsite4 and MCsite5 sites ranged from an average 0.61-152.28% and 9.17%-69.17%, respectively (Figure 2B-C). Pairwise comparisons indicated mutagenesis frequencies of MCsite4 sites can be categorized into three distinct groups (*p* value < 0.001): the high editing group (group H: MCsite4.8, 152.28%), the moderate editing frequency group (group M: MCsite4.4 with 17.04% and 4.7 with 16.20%), and the low editing frequency group (group L: MCsite4.1, 4.2, 4.3, 4.5, and 4.9, ranging from 0.61% to 1.71%) (Figure 2B). Comparison between the highest and lowest MCsite4 edited sites revealed a 249.64-fold difference. Similarly, mutagenesis frequencies at the MCsite5 sequences can also be grouped into the high, moderate and low editing groups (*p* value < 0.01) with 7.54-fold differences between the highest and lowest edited sites (Figure 2C). We then characterized the local sequence context surrounding each target site (25 bp from each side). High sequence similarities were found in these extended sequences for both MCsite4 and 5 families (Supplemental Figure S2B and C). Phylogenetic analyses within each target site family found no evident correlations between sequence similarity and the mutagenesis frequency groups (Supplemental Figure S2B and C). Thus, the local sequence context could not explain differential mutagenesis frequencies observed from individual target sites.

The initial association analysis of indel frequencies with DNA methylation domains and chromatin accessibility indicated unmethylated and accessible sites generally had higher mutagenesis levels than the methylated and inaccessible sites (Figure 2B-C). Notably, the strongly negative correlations between mutagenesis frequency and DNA methylation levels at both MCsite4 and 5 sites could also be observed when the cytosine methylation status was examined at the single nucleotide level of individual CRISPR target sites (Supplemental Figure S3). To systematically investigate the correlation between mutagenesis efficiency and chromatin features, we further characterized individual MCsites using all 23 chromatin features, including distinct histone modifications and histone variants (Supplemental Dataset 3) (Y. Liu et al. 2018). When hierarchical cluster analysis was performed, all lowly edited sites (group L) clustered with the heterochromatic features, such as H2A.W, H3K9me1, H3K9me2, H3K27me1, and hyper DNA methylation, for both MCsite4 and 5 (Figure 2D-E). On the contrary, the highly and moderately edited groups (group H and M) appeared to be associated with accessible, active chromatin features such as histone acetylation, H3K36me3, and H3K4 methylation (Figure 2D-E) (Roudier et al. 2011). To examine the impact of individual features, we performed correlation analyses by plotting mutagenesis efficiency at all 15 target sites with each chromatin and DNA methylation feature (Supplemental Figure S4). Consistent with the hierarchical cluster analysis, strong positive correlations were observed between mutagenesis frequency and euchromatin-related features such as H3K56ac (R = 0.85, *p* = 1.8e-13), H3K9ac (R = 0.82, *p* = 8.6e-12), H3K36ac (R = 0.75, *p* = 4.2e-9), H3K27ac (R = 0.71, *p* = 4.1e-8), H3K36me3 (R = 0.71, *p* = 3.5e-8), H3K4me3 (R = 0.65, *p* = 1.6–6), and accessibility (R = 0.51, *p* = 0.00038) measured with ATAC-Seq data (Supplemental Figure S4). On the contrary, strong negative correlations were found between mutagenesis frequencies and the heterochromatin-related features H2A.W (R = −0.57, *p* = 4.1e-5), H3K9me1 (R = −0.55, *p* = 0.00011), H3K9me2 (R = −0.55, *p* = 1e-4) and cytosine DNA methylation (R=-0.45, *p* = 0.0022) (Supplemental Figure S4).

### Improving CRISPR-Cas9 mutagenesis efficiency through combined reduction of DNA methylation

Because of the strong negative association observed between the lowly edited sites and heterochromatic features, we hypothesized that altering these chromatin states could improve mutagenesis efficiency at the refractory sites. In this study, we chose to perturb DNA methylation at the lowly edited sites because high levels of DNA methylation are often correlated with heterochromatic features (Crisp et al. 2020; Johnson et al. 2007; Zemach and Grafi 2003). We first sought to test the possible impact of CHG methylation on mutagenesis efficiencies because CHG methylation is often associated with H3K9me2 and heterochromatin (Springer and Schmitz 2017). To this end, the *chromomethylase3* (*cmt3-11t*) mutant was chosen due to the well-documented genome-wide reduction in CHG methylation (Stroud et al. 2013). Analysis of the single-base resolution DNA methylation profiles indicated that a substantial reduction in CHG methylation was confirmed at individual MCsite4 and 5 CRISPR targets in this mutant (Supplemental Figure S5). The CRISPR-Cas9 T-DNA constructs targeting MCsite4 and 5 sequences were transformed into the *cmt3-11t*mutant using the same procedure described above. As a result, transgenic *cmt3-11t* plants (T1) were identified with detectable mutagenesis activities at both MCsite4 and 5 target sites (Supplemental Figure S6). After normalizing to the CHLI2 control target site, we observed a nearly identical pattern in mutagenesis efficiency between WT and *cmt3-11t* plants for both MCsite4 and MCsite5 (Figure 3A-B). These results suggested that a reduction of CHG methylation alone is not sufficient to improve CRISPR-Cas9 mutagenesis at the refractory sites.

**Figure 3.**
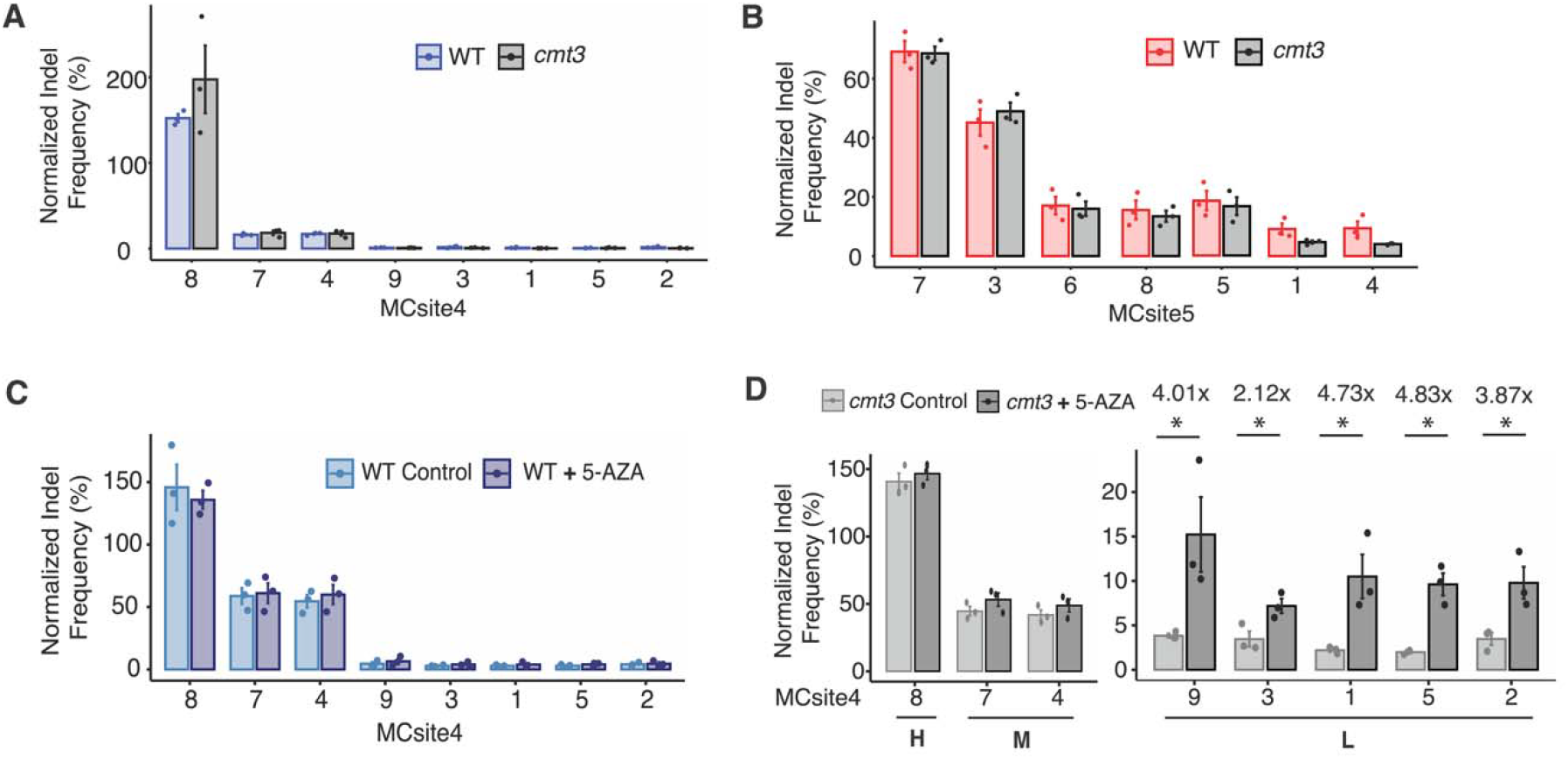
Comparisons of CRISPR-Cas9 mutagenesis frequencies using genetic and chemical approaches. *(A)* and *(B)* Normalized mutation frequency were compared for MCsite4 and MCsite5 sites in wild type and *cmt3* mutant background. *(C)* Normalized mutation frequencies for MCsite4 sites in wild type plants were compared with or without 5-azacytidine treatment. *(D)* Normalized mutagenesis frequencies were compared for the MCsite4 sites in the *cmt3* mutant plants with or without 5-azacytidine treatment. The MCsite4 sites were categorized in High (H; site 8), Moderate (M; site 7 and 4) and Low (L; sites 9, 3, 1, 5 and 2) mutagenesis groups. The standard error (SEM) was calculated for each target site with 3 replicates. The fold changes in mutagenesis were calculated for the sites in group L. Mann-Whitney U test was also performed with asterisks indicating significant *p*-values (*p* < 0.05).

Although additional genetic mutant lines with perturbed DNA methylation, such as *met1*, were available (Stroud et al. 2013), we have not been able to obtain transgenic CRISPR-Cas9 T-DNA events in these mutant lines. Therefore, we chose a chemical approach to reduce genome-wide DNA methylation in all 5mC contexts using 5-azacytidine (Griffin, Niederhuth, and Schmitz 2016). Analysis of the DNA methylation for 5-azacytidine treated plants revealed substantial reductions in DNA methylation at the highly methylated sequences at both MCsite4 and 5 (Supplemental Figure S7A) (Griffin, Niederhuth, and Schmitz 2016). We focused on MCsite4 in the subsequent 5-azacytidine experiment due to the lower mutagenesis frequencies at the lowly edited sites. The T2 plants with the CRISPR-Cas9 T-DNA targeting MCsite4 in the wildtype background were obtained from a self-pollinated T1 plant used above. After 2-weeks of growth with or without 100 μM 5-azacytidine, the plants were subjected to mutagenesis analyses using the NGS assay (Supplemental Figure S7B). No significant differences were observed for the normalized mutagenesis frequency at each MCsite4 site between the untreated and treated samples (Figure 3C). Thus, partial reductions in all DNA methylation contexts using 5-azacytidine did not reveal changes in mutagenesis frequencies.

Lastly, we tested the impact of combined reductions in DNA methylation using both the *cmt3-11t* mutant and 5-azacytidine treatments. T2 plants with the MCsite4-targeting T-DNA were obtained from a self-pollinated T1 plant in the *cmt3-11t* mutant background, grown for 2 weeks with or without 100 μM 5-azacytidine, and subjected to NGS analysis using the same procedure (Supplemental Figure S7C). After normalizing to the CHLl2 control, we observed two distinct patterns for mutagenesis efficiency across individual MCsite4 CRISPR targets. At the unmethylated and accessible sites (groups H and M), no significant differences in mutagenesis frequency were found between the 5-azacytidine treated mutants and the control group (Figure 3D). On the contrary, significant increases in mutagenesis frequency were observed at the inaccessible and hypermethylated sites (group L) when 5-azacytidine treatment was combined with the *cmt3-11t* mutant, *i*.*e*., MCsite4.9 (4.01-fold), MCsite4.3 (2.12-fold), MCsite4.1 (4.73-fold), MCsite4.5 (4.83-fold), and MCsite4.2 (3.87-fold) (Figure 3D). Together, these data indicated that strong reduction in DNA methylation in multiple contexts could result in significant improvement of CRISPR-Cas9 mutagenesis efficiency at the hypermethylated refractory sites.

### Differential CRISPR-Cas9 mutational profiles are associated with distinct chromatin features

CRISPR-Cas9 induced mutations are typically composed of either small deletions or 1 bp insertions (Allen et al. 2018). Small deletions are mainly derived from the NHEJ or MMEJ pathway through exonuclease-mediated end processing and ligation, while the 1 bp insertions were mostly generated from blunt-end or 1 bp staggered cleavage by Cas9 followed by DNA polymerase-mediated end filling (Gisler et al. 2019). To investigate the potential impact of chromatin context on CRISPR-Cas9 mutation outcome, we examined the insertion and deletion profiles for both MCsite4 and 5. As expected, most mutations contained 1 bp insertions and small deletions (< 10 bp) for both MCsite4 and 5 (Figure 4A). MCsite4 was preferentially repaired as 1 bp insertions, while MCsite5 showed a strong bias towards small deletion outcomes (Figure 4A). While the major mutation types at individual sites within each family seemed to be highly similar (Supplemental Figure S8A-D), further analysis revealed significant variations for the insertion rate between individual sites. The rate of insertion outcomes ranged from 56.25-81.73% and 7.57-30.27% in MCsite4 and MCsite5 sites, respectively (Figure 4B-C). We then conducted a correlation analysis using the 23 epigenetic features with the insertion rates at all 15 sites. Three histone H3 related features, H3K4me1 (R = −0.64, *p* = 0.01), H3.3 (R = −0.83, *p* = 3e-4), and H3.1 (R = −0.91, *p* = 2.8e-6), were identified with significant negative correlations with 1 bp insertional mutations (Figure. 4D; Supplemental Figure S9). Thus, these findings strongly suggested that chromatin features could not only affect CRISPR-Cas9 mutagenesis efficiency but also influence mutation outcomes.

**Figure 4.**
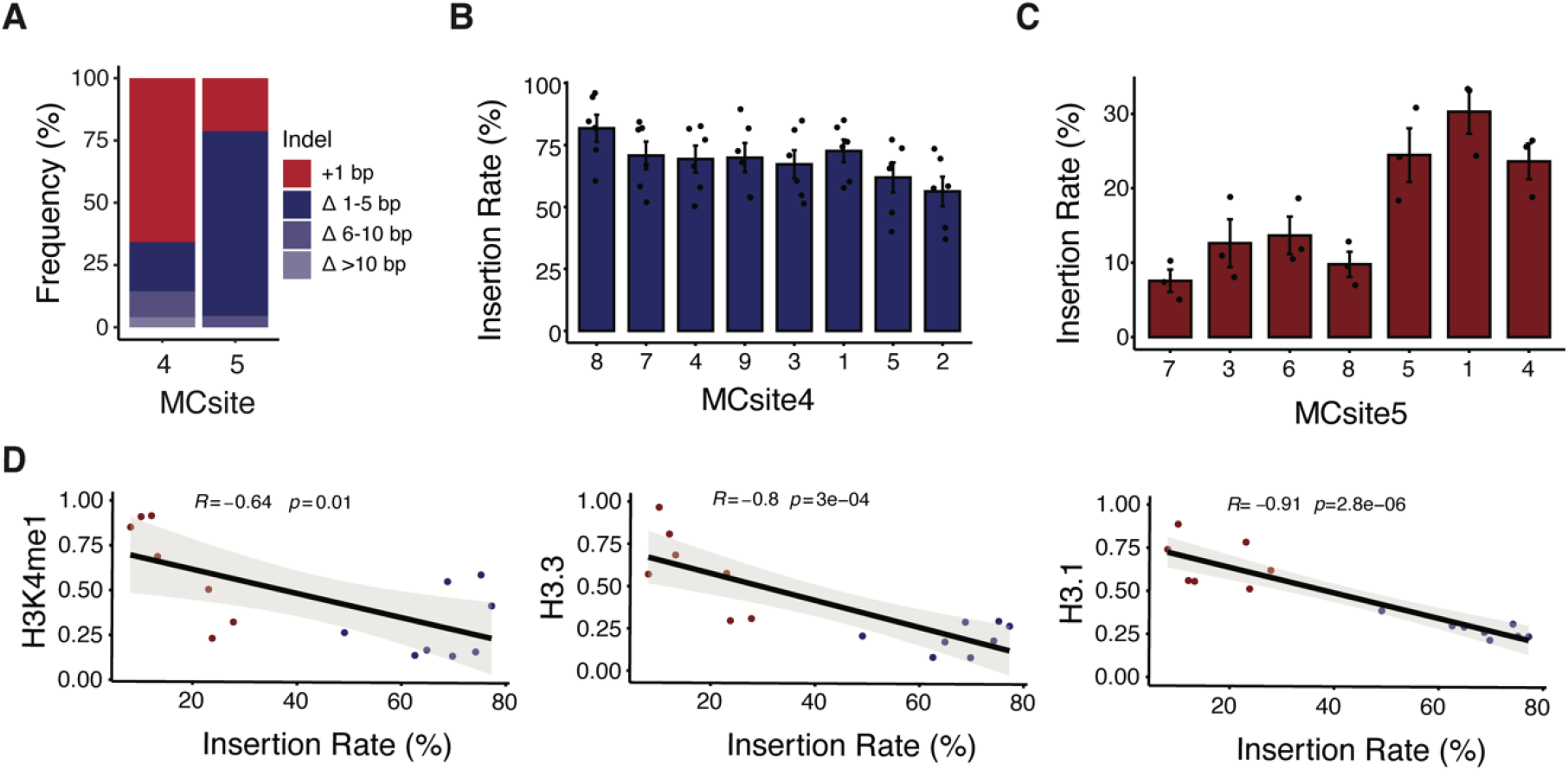
Analyses of CRISPR-Cas9 mutation profiles at MCsites. *(A)* Distribution of mutation outcomes at MCsite4 and MCsite5. The mutations were categorized as 1 bp insertions (+1 bp, red), 1-5 bp deletions (Δ 1-5 bp, dark blue), 6-10 bp deletions (Δ 6-10 bp, blue), and deletions of more than 10 bp (Δ >10 bp, light blue). *(B)* and *(C)* The 1 bp insertion frequencies at individual MCsite4 and MCsite5 targets. The standard error (SEM) was calculated for each target site with replicates (n=6 for MCsite4; n=3 for MCsite5). Kruskal-Wallis analysis indicated the p-value for MCsite4 and MCsite5 are 0.17 and 0.016, respectively. *(D)* Correlation plots between the insertion rate and 3 chromatin markers, H3K4me1, H3.3 and H3.1, with the blue dots for MCsite4 and the red dots for MCsite5. The trendline was plotted with gray indicating the standard error. The R value and *p*-values were calculated using Spearman’s rank correlation coefficient.

## Discussion

In recent studies, chromatin contexts have been demonstrated to have significant impacts on CRISPR-Cas9 mediated genome editing (Daer et al. 2017; G. Liu et al. 2019; Wu et al. 2014; Yarrington et al. 2018). Most of these findings indicated that heterochromatic features at the CRISPR target regions could impede CRISPR-Cas9 mutagenesis efficiency in multiple systems, such as yeast, rice, mouse, and human cell lines (Daer et al. 2017; G. Liu et al. 2019; Wu et al. 2014; Yarrington et al. 2018). However, it remained unclear whether heterochromatic features could only temporarily delay CRISPR-Cas9 mutagenesis (Kallimasioti-Pazi et al. 2018). In this study, we systematically characterized the impact of 23 distinct DNA methylation and chromatin features on CRISPR-Cas9 mutagenesis efficiency and mutation outcomes by investigating CRISPR-Cas9 transgenic Arabidopsis plants. Consistent with the previous studies, our results demonstrated that inaccessible and heterochromatic features were strongly associated with low mutagenesis efficiency. Such repressive effects can be long-lasting through plant development leading up to a 250-fold difference between identical CRISPR target sites. On the other hand, mutagenesis efficiency at the target sequences in accessible chromatin regions could also have significant variations ranging from 1.53-fold to 10-fold at MCsite5 and MCsite4, respectively (Figure 2B-C). This observation suggested that specific chromatin features other than just open or closed chromatin should be considered to account for CRISPR-Cas9 mutagenesis efficacy. Close examination of individual chromatin features identified several euchromatic marks, such as H3K9ac, H3K56ac, H3K36ac, H3K27ac, H3K4 methylation and H3K36m3, that were positively correlated with mutagenesis efficiency. Further investigation with a larger data set will be needed to dissect their impacts on CRISPR-Cas9 mutagenesis in greater detail.

In this study, we observed strongly negative correlations between CRISPR-Cas9 mutagenesis efficiency and repressive chromatin features, such as hyper DNA methylation, low DNA accessibility, H3K9 methylations. To test the hypothesis that modulating some of these features could improve mutagenesis frequency at the lowly edited sites, we sought to reduce DNA methylation by using both genetic and chemical approaches. Our results indicated that partial reduction of DNA methylation using either a mutant affecting CHG methylation or 5-azacytidine treatment alone was not sufficient to improve mutagenesis efficiency at any of the tested sites. When combining the CHG deficiency mutant with 5-azacytidine chemical treatment, 2.1 to 4.8-fold improvements in mutagenesis efficiency were found at the lowly edited sites but not at the highly and moderately edited sites. Combined reduction of DNA methylation in multiple contexts has been demonstrated to increase chromatin accessibility and even alter the higher order 3D chromatin organization in Arabidopsis (Zhong et al. 2021). Thus, mutagenesis efficiency improvement at lowly edited sites observed here could have resulted from increasing chromatin accessibility and/or changing 3D chromatin organization due to the significant reduction of DNA methylation at multiple contexts.

In addition to the impacts on mutagenesis efficiency, chromatin features have been suggested to influence CRISPR-Cas9 mutation outcomes. For example, a recent study with more than 1,000 copies of identical insertion sites indicated the 1 bp insertions were found more prevalent in euchromatin than in heterochromatin, likely through recruiting different DNA repair machinery (Schep et al. 2021). However, conflicting results were also reported showing little impact of chromatin features on mutation outcomes (Gisler et al. 2019; Kallimasioti-Pazi et al. 2018). In this study, we observed significant variations for the 1 bp insertion rate in different chromatin contexts. Of the 23 chromatin features analyzed, three distinct histone H3 features, H3K4me1, H3.3, and H3.1, exhibited significantly strong negative correlations with the 1 bp insertion rate. Interestingly, H3K4me1 was also identified by Schep et al. as a marker to correlate with distinct mutation outcomes, while H3.1 and H3.3 were not included in their study (Schep et al. 2021). This suggested that the balance between 1 bp insertions and small deletions could be influenced by specific chromatin markers. Distinct chromatin features have been reported to recruit different DNA repair machinery and result in different repair outcomes in mammalian systems (Fnu et al. 2011; Jacquet et al. 2016; Luijsterburg et al. 2016). Further investigation is needed to address the potential roles of specific chromatin markers in determining DNA repair outcomes.

We proposed a model to account for the impacts of chromatin features on CRISPR-Cas9 mutagenesis efficiency and mutation outcomes (Figure 5). In the first step, chromatin features are the key determinants for CRISPR-Cas9 recognition and binding efficiency (Figure 5). In general, heterochromatic features, such as H3K9 methylations, H3K27me1, H2A.W, and hyper DNA methylation, could significantly reduce chromatin accessibility and thus reduce the recognition and binding efficiency of CRISPR-Cas9. Such repressive effects could persist through cell division and development. After CRISPR-Cas9 locates the target site and binds the genomic DNA, the Cas9 nuclease can introduce double stranded DNA breaks either with a 1-bp 5’ overhang (staggered cut), or blunt ends (Gisler et al. 2019). The cleaved product can be repaired to yield three outcomes: wild-type sequence by perfect ligation, 1 bp insertions, or small deletions (less than 10 bp) (Figure 5). It has been proposed that the staggered cut primarily leads to 1 bp insertion via template-dependent repair, facilitated by a DNA polymerase, while the blunt cut mainly results in small deletions through end resection (Schmid-Burgk et al. 2020; Gisler et al. 2019). The blunt ends could occasionally be repaired by DNA polymerase, likely the members from DNA polymerase family X, without DSB end resection, resulting in template-independent 1 bp insertions (Gisler et al. 2019). In this study, we observed both templated 1 bp insertion and template-independent insertions, as exemplified in MCsite5 and MCsite4, respectively (Supplemental Figure S8A-B). Nevertheless, the balance between 1 bp insertion and small deletion products is primarily dependent on the repair choices between short-range DSB end resection and DNA polymerase end filling (Figure 5) (Schmid-Burgk et al. 2020; Lemos et al. 2018). While sequence features are a key determinant for mutation profiles, our data indicated that chromatin features, such as H3K4me1, H3.3, and H3.1, could also significantly impact the balance between 1 bp insertion and small deletion outcomes. These specific chromatin features may exert their influences on the balance between the staggered and blunt cut, or through modulating the balance between DNA polymerase end filling and short-range DSB end resection during NHEJ. In fact, previous studies have demonstrated that distinct chromatin features, such as the H3.3 variant, could impact DSB end resection (Luijsterburg et al. 2016).

**Figure 5.**
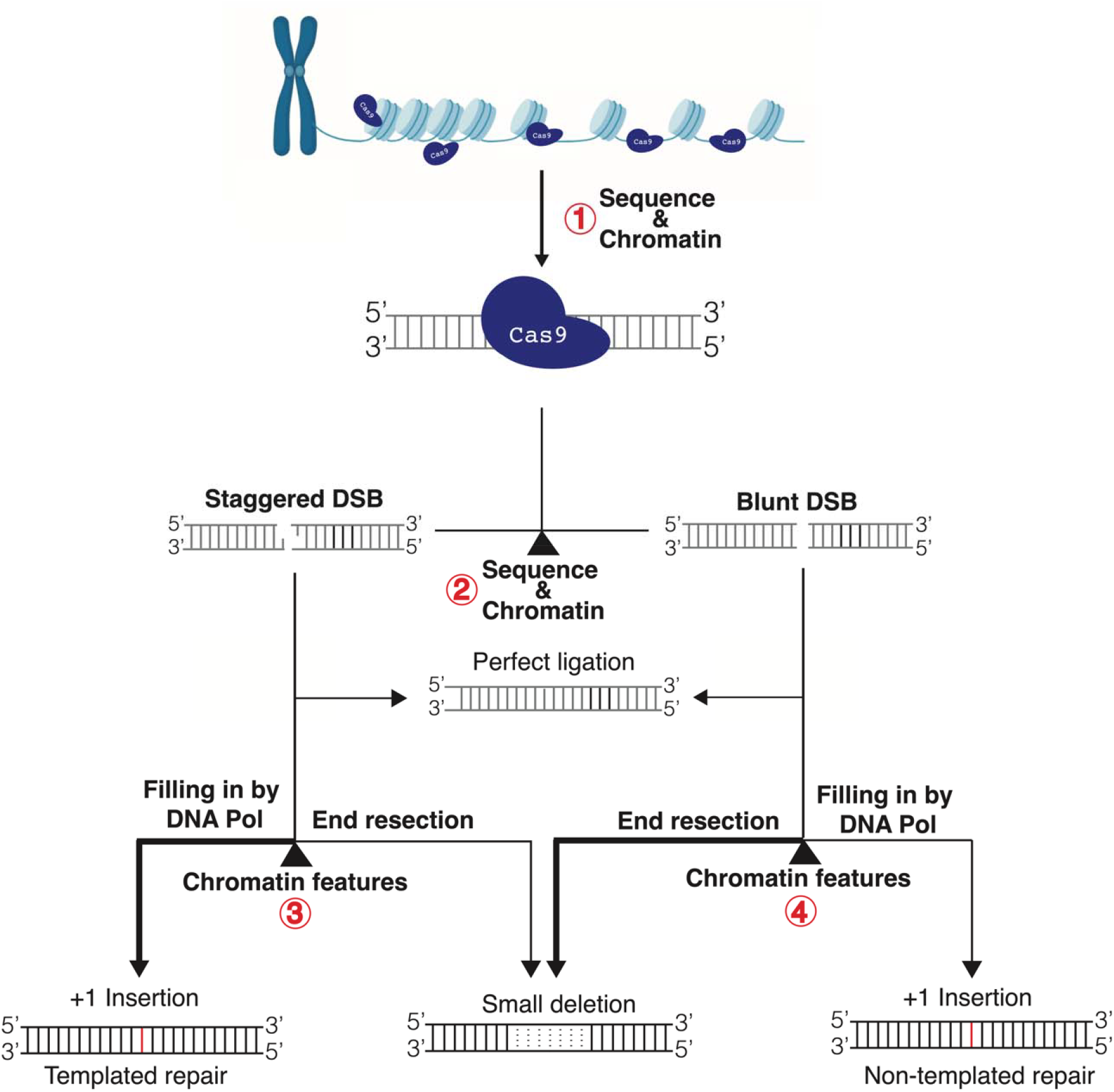
Model depicting the factors that influence CRISPR-Cas9 mutagenesis efficiency and mutation outcomes. The four key check points were indicated in the numerical order with red circles. At Point 1, both sequence and chromatin features are the key determinants for CRISPR-Cas9 recognition and binding efficiency. At Point 2, sequence and chromatin features are crucial to determine staggered or blunt cleavage by CRISPR-Cas9. The cleaved products can be ligated back perfectly or go to Point 3 or 4, where chromatin features likely influence occurrence of end resection. With end resection, it generates small deletions. When the end resection is limited, the broken ends will be directly filled by DNA polymerase (DNA Pol) to generate 1 bp insertions. 1 bp insertions can be the products from template-dependent repair using the 1 bp 5’ overhang by staggered cleavage, or from template-independent repair by filling in blunt ends. The staggered cleavage primarily leads to 1 bp insertions, while the blunt cut mainly results in small deletions as indicated by line thickness.

Recently much effort has been made to develop computational tools to predict CRISPR-Cas9 editing efficiency and/or repair outcomes. To our knowledge, these tools primarily relied on sequence features (Allen et al. 2018; Concordet and Haeussler 2018; Xiang et al. 2021). Yet our results indicated that non-sequence features should also be taken into consideration for the prediction of both CRISPR-Cas9 efficiency and mutation outcomes. One implication from this study is that heterochromatin related features, such as low accessibility, H3K9me1, H3K9me2, H3K27me1, H2A.W and hyper DNA methylation, should be avoided in order to design gRNAs with high efficiency. When it comes to predicting CRISPR-Cas9 mutation profiles, chromatin features such as H3K4me1, H3.3, and H3.1 should also be considered. Future studies using genetic mutants involved in histone medications will be interesting to further dissect the impact of distinct chromatin features on CRISPR-Cas9 mutagenesis. A better understanding of the interplay between chromatin dynamics and CRISPR-Cas9 will enable the development of more precise and efficient genome engineering technologies.

## Methods and Materials

### Identification of multicopy CRISPR sites

To identify gRNAs that matched to multiple places in the genome, the Arabidopsis Col-0 reference genome (TAIR10) was parsed to identify every NGG PAM site using *seqkit locate* (Shen et al. 2016) with the search motif NNNNNNNNNNNNNNNNNNNNNGG. The number of occurrences of each non-redundant sequence in the genome was summarized using *csvtk*. The resulting distinct target sites were then filtered in R (v4.1.2) to retain gRNAs that had between 7 to 25 distinct matches. Sites were then eliminated if they had simple sequence motifs consisting of 5 As, Ts, Gs or Cs in a row; CG content outside 40%-60% or if the sequence was found in the chloroplast or mitochondria genome; to give a final list of 9,902 candidate gRNAs. The gRNA identification script is available at https://github.com/pedrocrisp/Weiss_et_al_gRNA_chromatin. The potential gRNAs were also screened to identify those with restriction enzyme recognition motifs (from a list commercially available enzymes from NEB) that overlapped position 17-18 of the gRNA (between position 3 and 4 bp from the PAM) such that an indel mutation in this position would disrupt restriction enzyme recognition and cleave for efficient screening of edited transgenic lines. A final list of 7,971 gRNAs were then annotated with chromatin state information by overlapping the coordinates of the gRNA target sites with chromatin annotation files using *bedtools (Quinlan and Hall 2010)*. Chromatin data included nine histone states (Sequeira-Mendes et al. 2014); chromatin accessibility (Lu et al. 2016); DNA methylation (Crisp et al. 2017; Springer and Schmitz 2017). DNA methylation data was converted to methylation domains for each 100 bp non-overlapping window of the TAIR10 genome using the method detailed in (Crisp et al. 2020). Chromatin accessibility at the CRISPR target site was called either Open or Closed based on the presence of an overlapping ATAC-Seq peak or lack thereof, respectively, using the accessibility profiles in (Lu et al. 2016). Gene and transposable element annotations were downloaded from Araport v11.

### T-DNA vector construction

The CRISPR-Cas9 constructs were created using the Golden Gate assembly method. The gRNA sequences were first assembled into the pMOD_B2301 backbone containing the gRNA targeting the MCsite and the CHLl2 positive control (Čermák et al. 2017). T-DNA constructs were assembled using pMOD_A0101 (Cas9), pMOD_B2301 containing the gRNA array, pMZ105 (luciferase reporter), and pTRANS230d (T-DNA backbone) (Čermák et al. 2017). All of the T-DNA constructs described in this study are available at Addgene. The modular components used to build the T-DNA plasmids can be found at https://www.addgene.org/browse/article/28189956/ (Čermák et al. 2017).

### Plant materials and growth conditions

The *Arabidopsis thaliana* Columbia ecotype (Col-0) was used in these experiments. The *cmt3-11t* (stock CS16392) genotype was acquired from the Arabidopsis Biological Resource Center (ABRC). Floral dip transformation was performed according to the protocol as previously outlined (Zhang et al. 2006). Transgenic T1 seeds were sown on soil and exposed to BASTA selection to recover transgenic plants. Plants were grown in a growth chamber with the following conditions: 16/8 hours light/dark cycle, 22 °C, and 55% humidity.

### Characterization of chromatin features

Previously analyzed datasets (BigWig files) for the 23 chromatin features were downloaded from Plant Chromatin State Database (PCSD) (Y. Liu et al. 2018). The datasets for wild type Arabidopsis with and without 100 μM 5-azacytidine treatment was downloaded from Griffin et al. (Griffin, Niederhuth, and Schmitz 2016). For each dataset, the values for the 1kb window (500 bp upstream and 500 bp downstream from the center of gRNA) were calculated using the Deeptools2 computeMatrix in the reference-point mode. Similarly, the values for each nucleotide of target sites were calculated using the scale-region mode at single base resolution. Each dataset was then normalized to allow for comparison on a scale of 0-1 with 1 indicating the highest level of that feature.

### 5-azacytidine treatment and luciferase screening

T2 seedlings from self-pollinated T1 Arabidopsis plants were grown on 1% agar plates containing 0.5 Murashige and Skoog (PhytoTech Labs) and 100 μM 5-azacytidine (Griffin, Niederhuth, and Schmitz 2016). After two weeks, seedlings were screened for presence of the transgene using a luciferase reporter. The luciferase assay procedure was performed using the Bio-Glo™Luciferase Assay System (Promega Corp., Madison, WI, USA) in accordance with the manufacturer’s instructions.

### Mutation genotyping and characterization of mutation profiles

Genotyping was performed using two methods: genomic PCR followed by restriction enzyme digestion (CAPS) and the NGS assay using Illumina paired-end read amplicon sequencing. All tissues for genotyping were collected at two weeks post germination for the CTAB-base genomic DNA extraction. PCR was performed using GoTaq Green Mastermix (Promega Corp., Madison, WI, USA) according to the manufacturer’s instructions, with an annealing temperature of 55 °C (CHlL2 and MCsite4) or 60 °C (MCsite5) with an extension time of 1 minute. Primers to amplify CHLl2, MCsite4, and MCsite5 target sites can be found in Table S1. Amplicons were then subjected to restriction enzyme digestion using *BsmA*I (CHlL2), *Alu*I (MCsite4), or *Drd*I (MCsite5) according to the manufacturer’s instructions. PCR amplicons generated with the corresponding primers were subjected to Illumina paired-end read sequencing (Genewiz Inc., South Plainfield, NJ, USA). The raw NGS reads were analyzed using CRISPResso2 to estimate indel mutation rates (Clement et al. 2019). To analyze the mutation. To analyze the mutation profiles for each sample, the NGS reads with indel mutations were extracted from the CRISPResso2 output files with a 2% threshold. The resulting output files were then loaded into R studio (version 4.1.0) for data visualization using ggplot (41). The total read counts for CRISPR-Cas9 editing frequency and repair profiles can be found in Table S2. Normalized indel frequencies were calculated by dividing the indel frequency of each MCsite by the CHlL2 positive control indel frequency within each replicate.

## Data availability

All sequencing data analyzed in this manuscript will be available at the National Center for Biotechnology Information (NCBI) under BioProject Accession PRJNA795172 (Supplemental Dataset 4).

## Acknowledgements and Funding

We thank all members of the Zhang Lab for helpful discussion. M.S. was supported by the Undergraduate Research Opportunities Program (UROP) from the University of Minnesota. Funding from NSF IOS-1934384 to N.M.S. partially supported this work. P.A.C. was supported by an ARC Discovery Early Career Award (DE200101748). F.Z. was supported by the startup fund from Department of Plant and Microbial Biology, University of Minnesota.

## Author Contributions

T.W., P.A.C., N.M.S. and F.Z. designed research; T.W., P.A.C., K.M.R., and M.S. conducted experiments; T.W., K.M.R., and F.Z. analyzed data; T.W., N.M.S. and F.Z. wrote the paper.

## Conflict of Interest

The authors declare no conflict of interest.

## Notes

### Competing Interest Statement

The authors have declared no competing interest.

### Summary of Updates

Supplemental files uploaded

