## Supplemental Information for "Drastic differential CRISPR-Cas9 induced mutagenesis influenced by DNA methylation and chromatin features"

**This file includes:**

Figures S1 to S9

Tables S1 to S2

Legends for Datasets S1 to S3

SI References

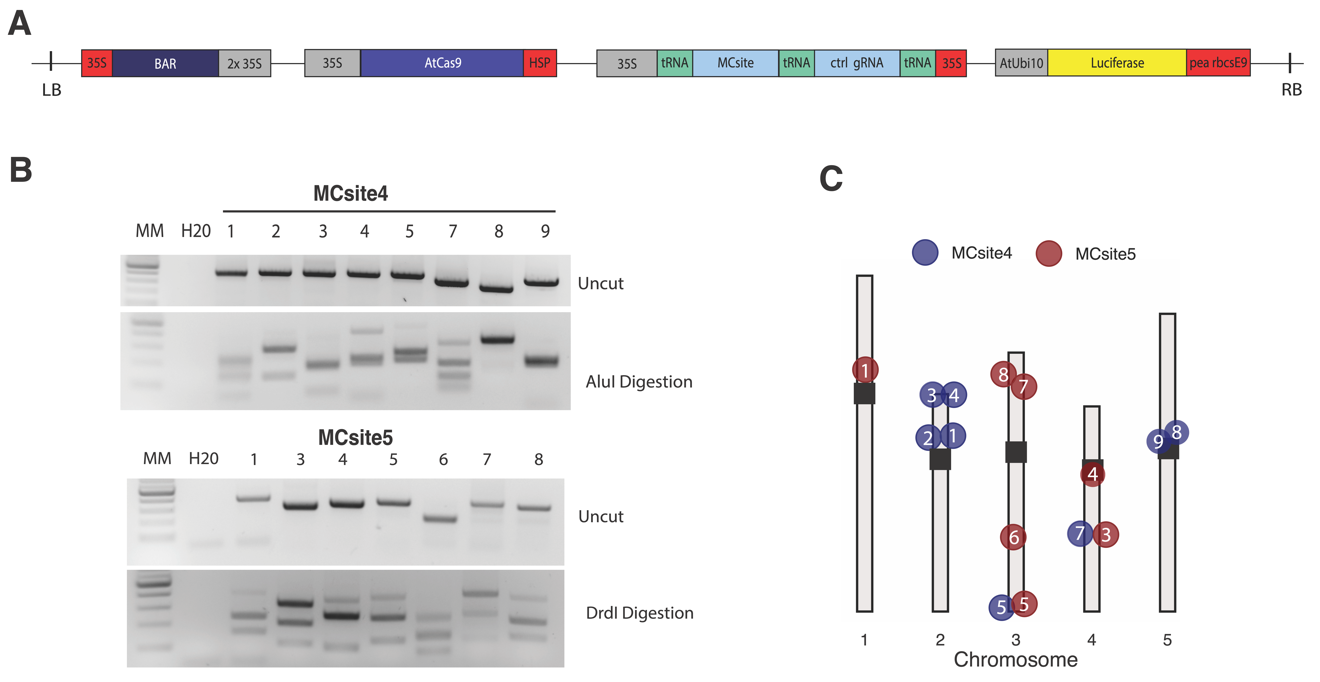

Fig. S1. Characterization of MCsites for CRISPR-Cas9 mutagenesis. (A) Illustration of the T-DNA constructs. (B) Representative CAPS genotyping images for MCsite4 and MCsite5. Samples were genotyped by genomic PCR (Uncut) and the CAPS assay (Alu Digestion for MCsite4 and DrdI digestion for MCsite5), with a 1-kb ladder, and no genomic DNA control (H20). (C) Distribution of the CRISPR target sites of MCsite4 (blue) and MCsite5 (red). Gray bars represent each chromosome with the black box indicating the centromere. The white number inside of each colored circle corresponds to the CRISPR target for that MCsite. Site 6 from MCSite4 and Site 2 from MCSite5 were not amplifiable with the site-specific PCR primers. Thus, they were excluded from the CAPS and NGS assays.

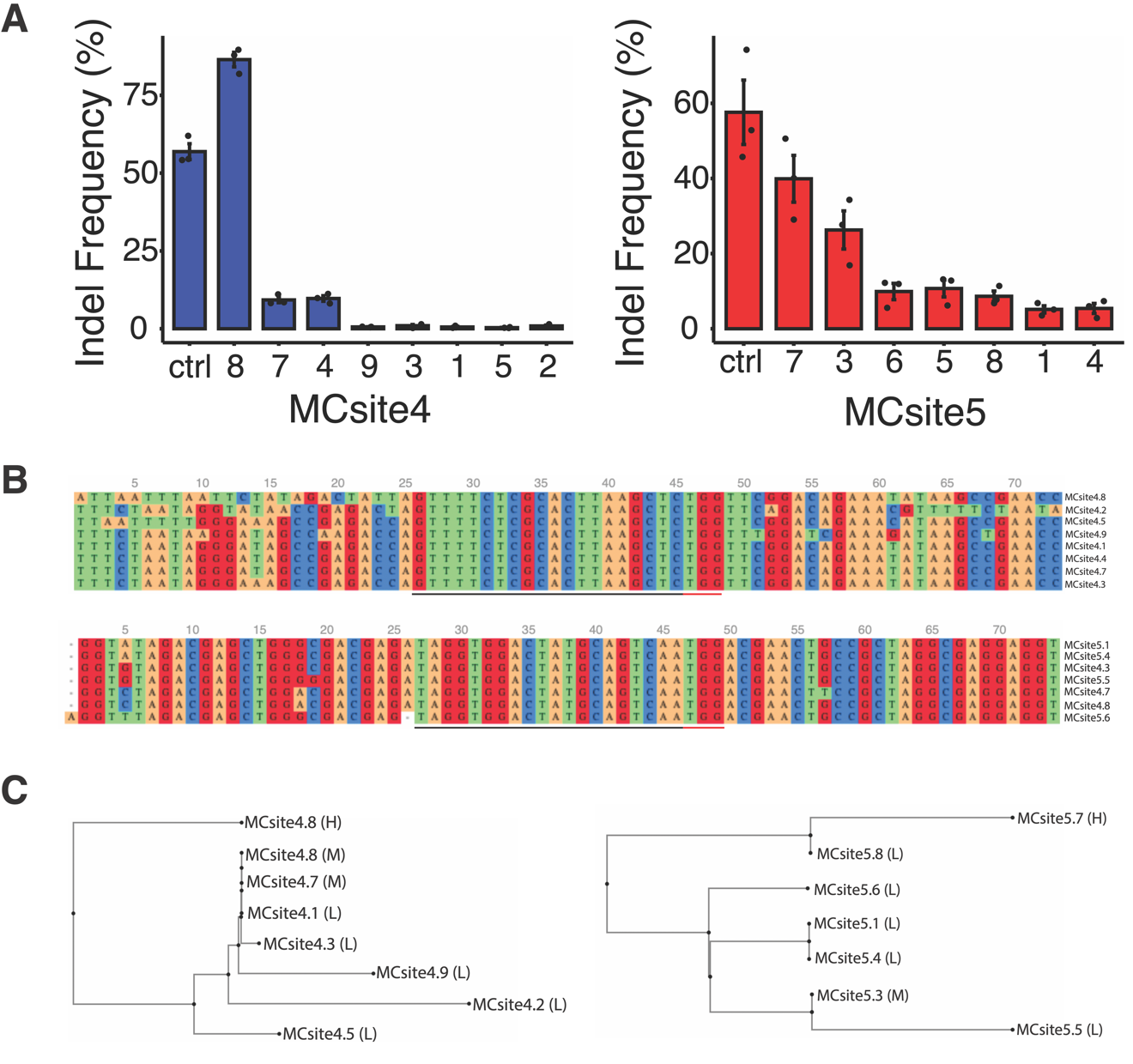

Fig. S2. Non-normalized mutagenesis efficiency and sequence comparison for individual target sites in MCsite4 and MCsite5. (A) Bar graphs displaying the non-normalized mutagenesis frequencies at CRISPR targeted sites for MCsite4 (blue) and MCsite5 (red). Ctrl is abbreviated for the CHLl2 site.  The standard error (SEM) is displayed for each target site with the dots indicating independent replicates (n = 3). (B) Sequence alignments of the CRISPR targets for MCsite4 and MCsite5. The sequence includes the 25 nucleotides to the left and to the right of the protospacer (underlined by a black bar) and PAM sequence (underlined by a red bar). The dendrogram indicates similarity between the sequences. Alignment was created using the MAFFT version 7 online tool with default settings (https://mafft.cbrc.jp/alignment/server/) (1). (C) Sequence similarity dendrograms were created using the Neighbor-Joining (NJ) method on MAFFT version 7 online tool with default settings (https://mafft.cbrc.jp/alignment/server/). H (high), M (moderate), and L (low) next to each target site indicate the mutagenesis group that CRISPR site is associated with.

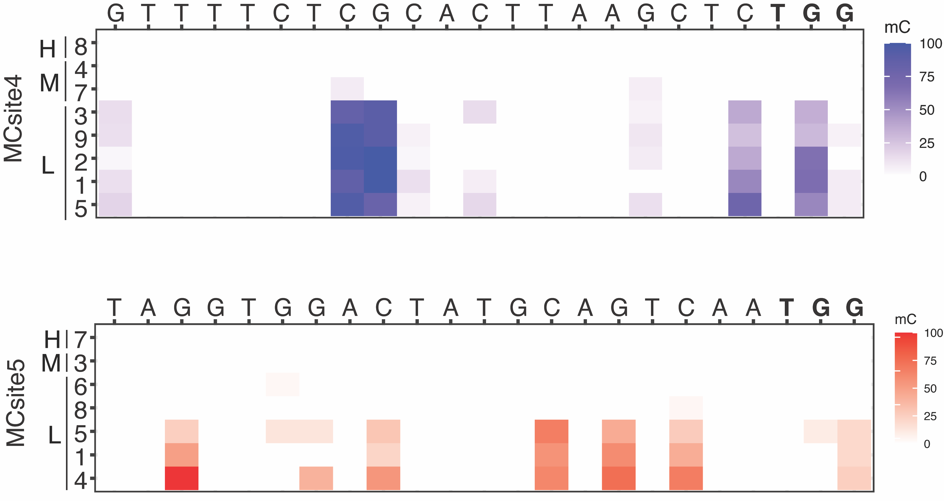

Fig. S3. Single nucleotide heatmap of DNA methylation levels at MCsite4 (blue) and MCsite5 (red) protospacer and PAM sequences (red) ranging from 0 (unmethylated) to 100 (fully methylated). H (high), M (moderate), and L (low) along the y-axis indicate which mutagenesis group that CRISPR site is associated with. The PAM sequence is bold.

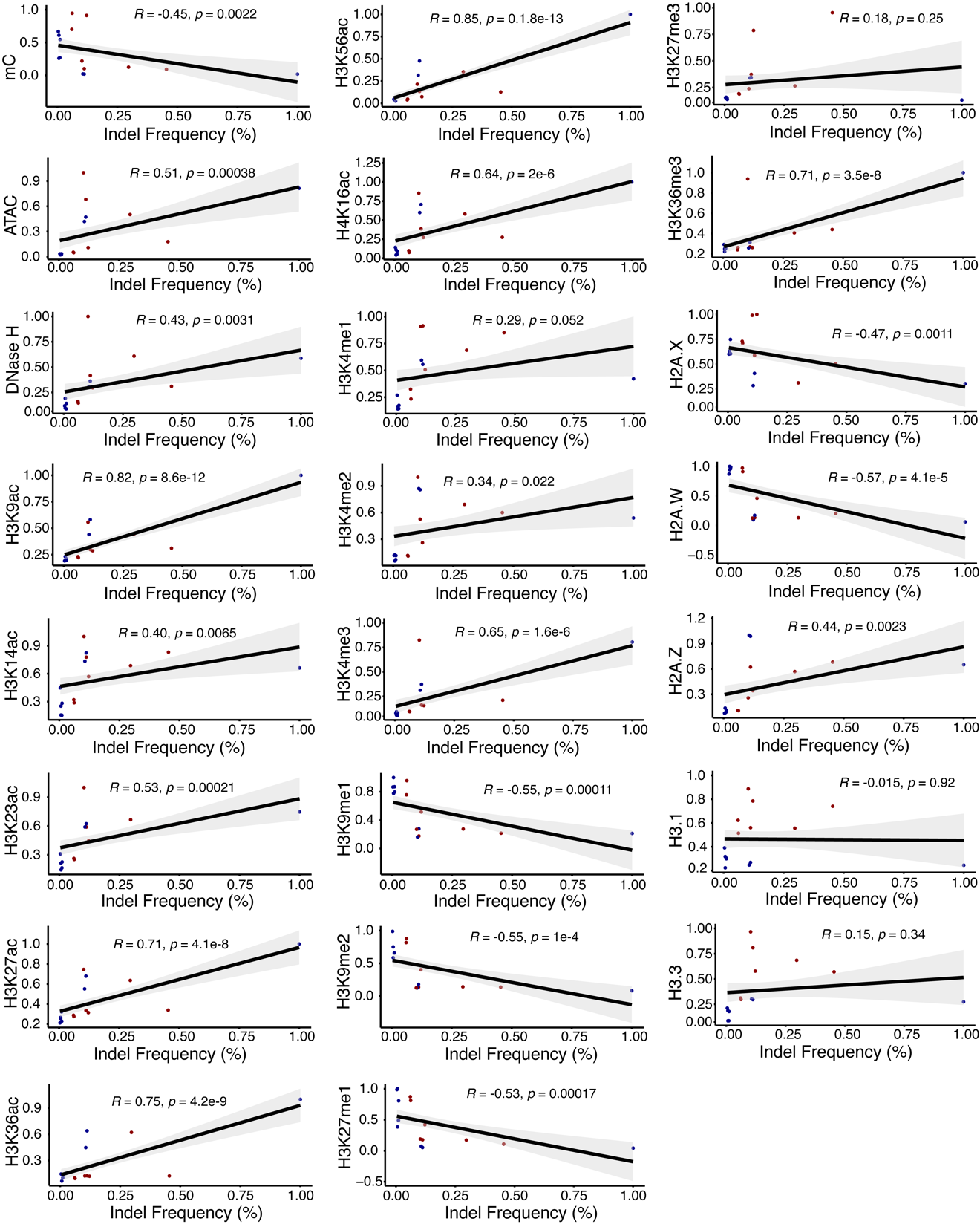

**Fig. S4.** Correlation analysis for CRISPR-Cas9 mutagenesis frequencies and chromatin features. The blue dots represent MCsite4 and the red dots represent MCsite5. The trendline is black with gray indicating the standard error. The R value and p-values are indicated at the top of each correlation plot according to Spearman’s rank correlation coefficient. Each feature was normalized using all 15 target sites on a scale of 0 to 1, with higher values indicating the higher levels for the respective feature.

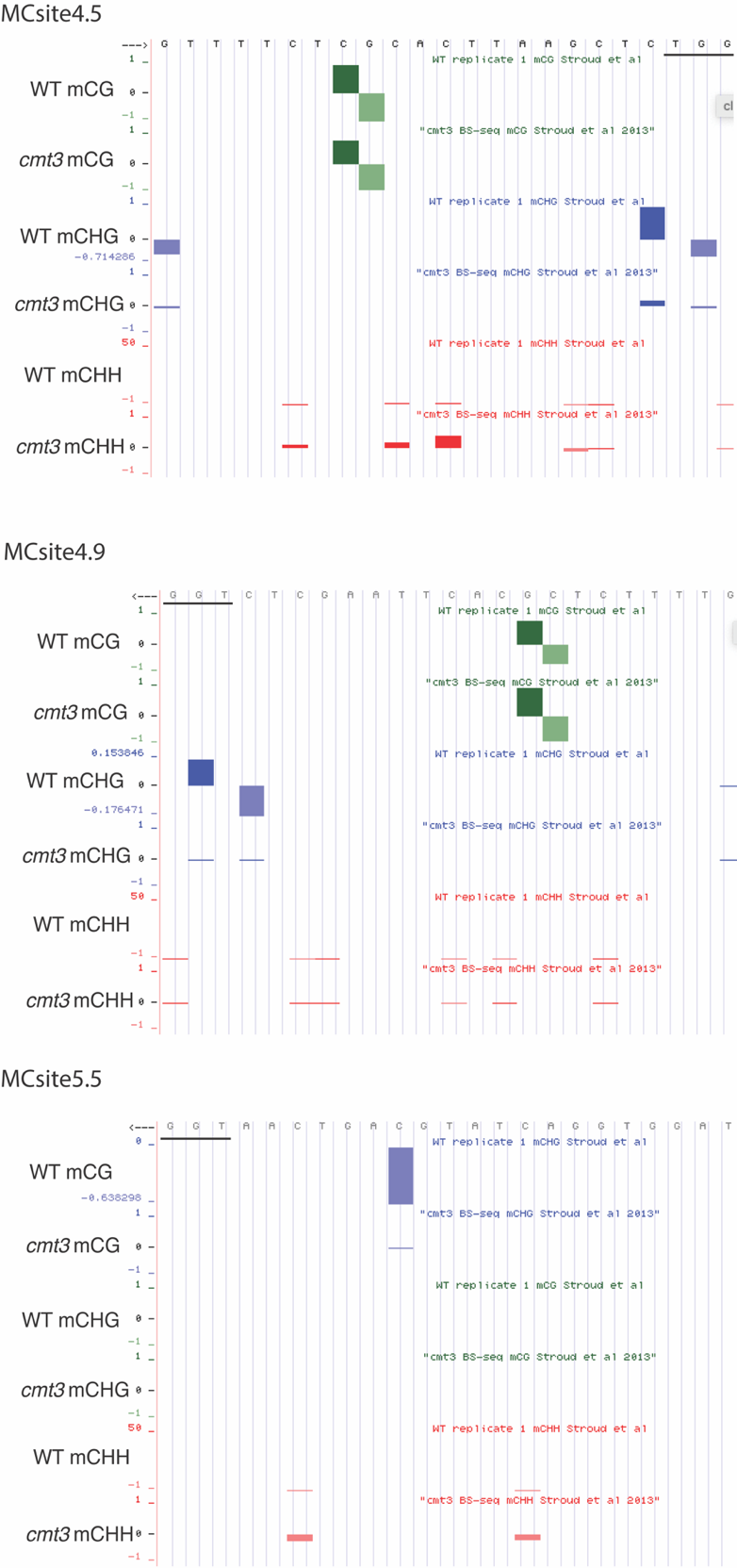

**Fig. S5.** Characterization of the single-based DNA methylation status at MCsite4 and MCsite5 in the wild type and *cmt3* mutant plants. Representative screenshots of MCsite4.5, MCsite4.9, and MCsite5.5 from the UCSC genome browser (2). The protospacer sequence is displayed along the top of each screenshot with the PAM underlined. 6 individual methylome tracks (wild type mCG context, cmt3 mCG context, WT mCHG context, cmt3 mCHG context, WT mCHH context, and cmt3 mCHH context) displaying the levels of DNA methylation at each individual nucleotide along the protospacer and PAM genomic DNA sequence. Green corresponds with mCG, blue with mCHG and red with mCHH.

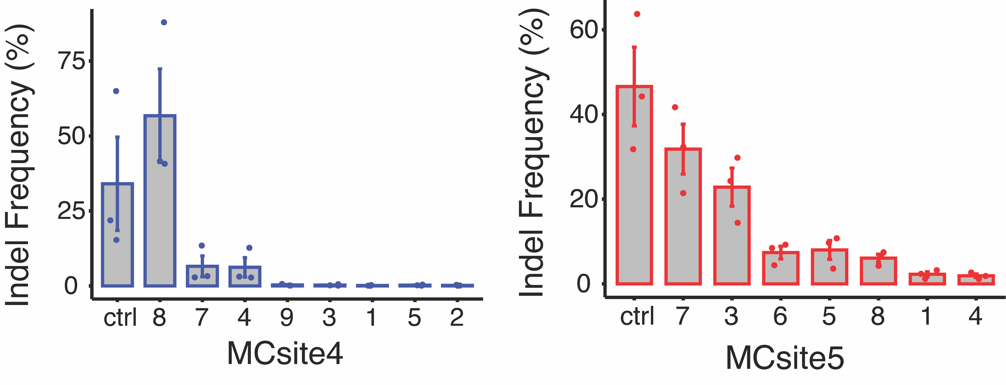

**Fig. S6.** Unnormalized mutagenesis frequencies for MCsite4 (blue and gray) and MCsite5 (red and gray) in the *cmt3* mutant plants. Ctrl is abbreviated for the CHLl2 site. The standard error (SEM) is displayed for each target site with the dots indicating independent replicates (n = 3).

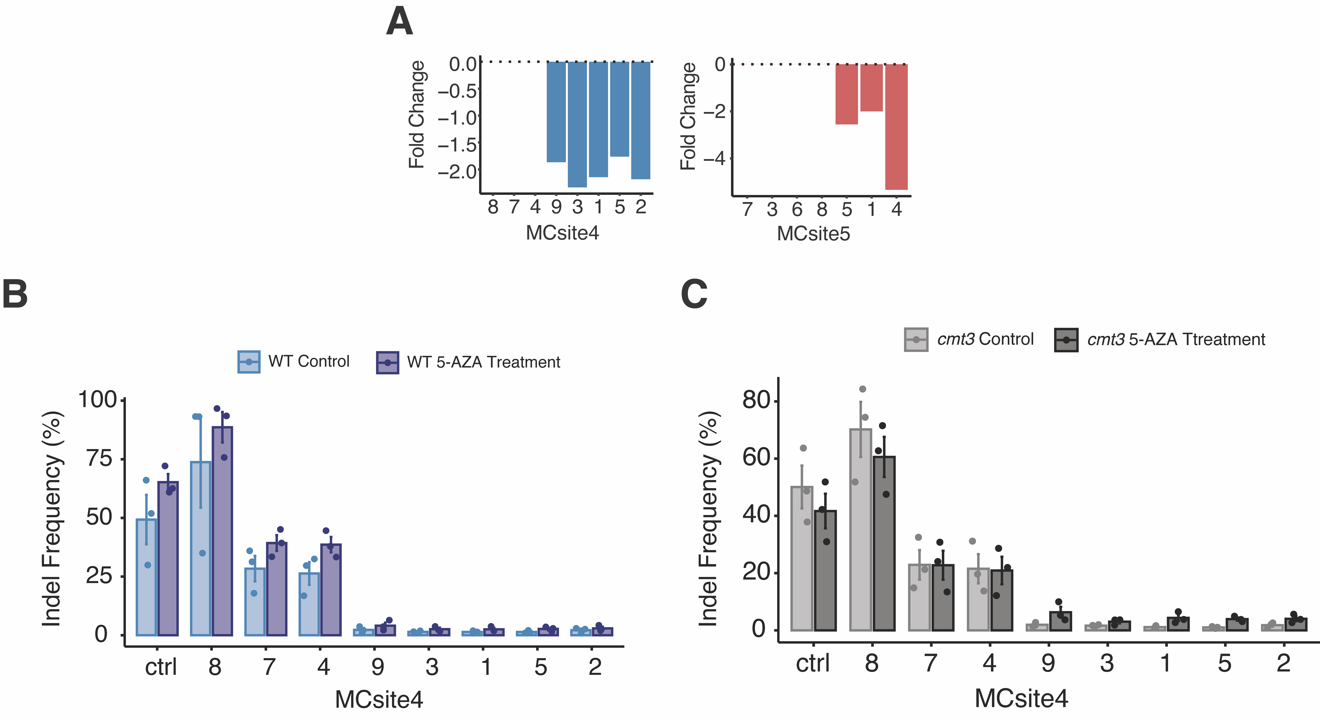

**Fig. S7.** 5-azacytidine treatment of the wild type and *cmt3* T2 seedlings. (A) DNA methylation reduction fold changes in a 1kb window (500 bp upstream and downstream from the CRISPR-Cas9 cut site) in the 100μM treated 5-azacytidine samples relative to the mock untreated samples from (3). Fold change was calculated by dividing the WT mock nontreated value by the 5-AZA 100 μM value, and then multiplied by -1. The dotted line at the top of each bar graph represents zero change, and each bar is color coded as either blue (MCsite4) or red (MCsite5). (B) and (C) Unnormalized mutagenesis frequencies for MCsite4 and the CHlL2 control (ctrl) in the wild type plants with and without 5-azacytidine treatment (B), and in the cmt3 mutant plants with and without 5-azacytidine treatment (C). The standard error (SEM) is displayed for each target site with the dots indicating replicates (n = 3).

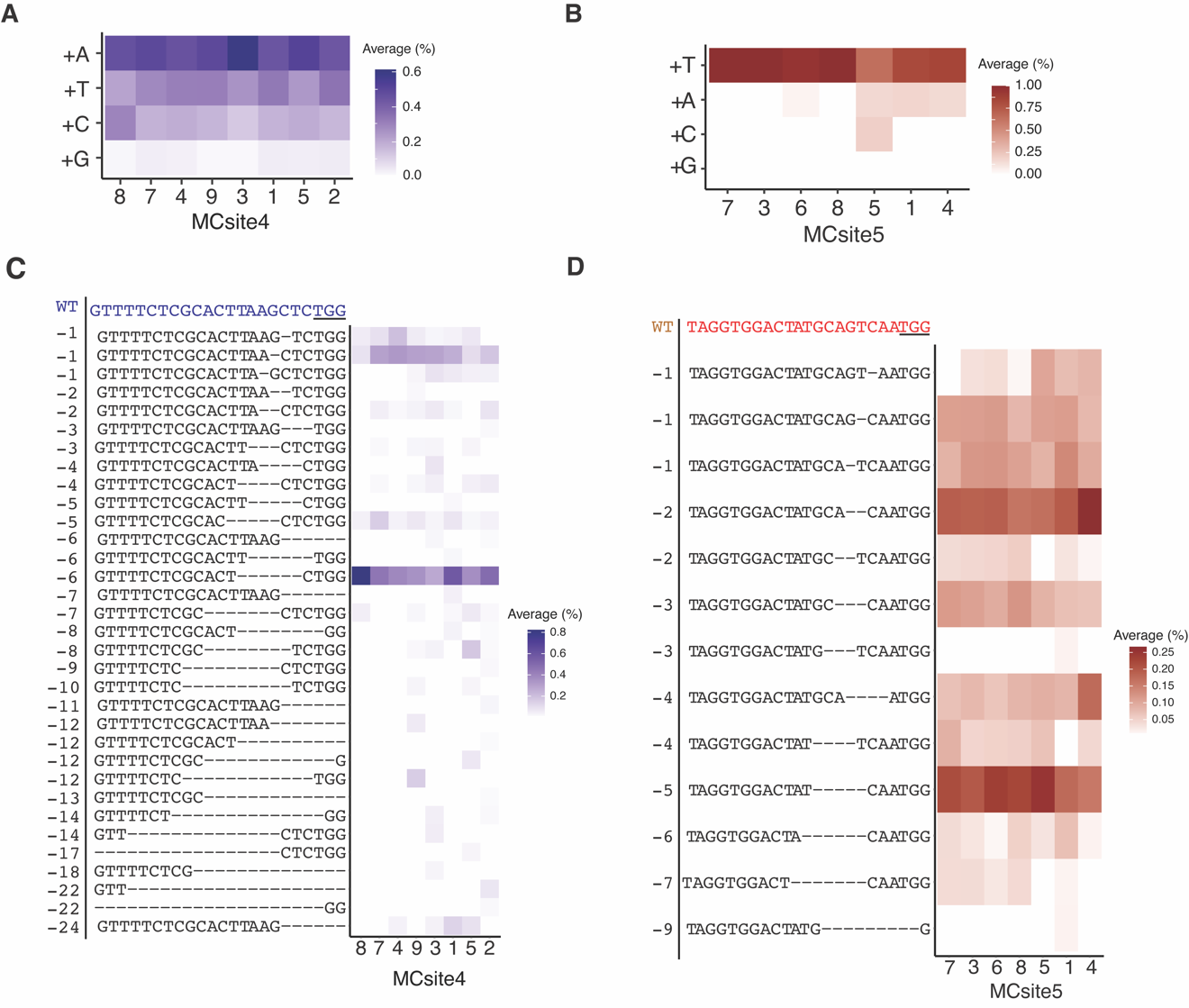

**Fig. S8.** Characterization of mutation outcomes for MCsite4 (blue) and MCsite5 (red). (A) and (B) Heatmap displaying the frequency of 1 bp insertions that occurred at each MCsite4 (A) and MCsite5 (B) sites in wild type plants. In the MCsite4 sites, the majority of 1 bp insertions (A, T or C) was derived from template-independent DNA polymerase-mediated end filling. In the MCsite5 sites, the majority of 1 bp insertions (T) was derived from templated-dependent end filling. (C) and (D) Heatmap displaying the frequency of deletion outcome that occurred at each MCsite4 (C) and MCSite5 (D) sites in wild type plants. The wild type sequence is at the top of the y-axis with the PAM underlined. The number to the left of each deletion outcome indicates the size of the deletion. The frequency of each repair outcome was calculated by using the total number of reads with insertions or deletions divided by the total number of mutated reads for each site. This was done for all three replicates. The average of the three replicates was then calculated and plotted as a heatmap.

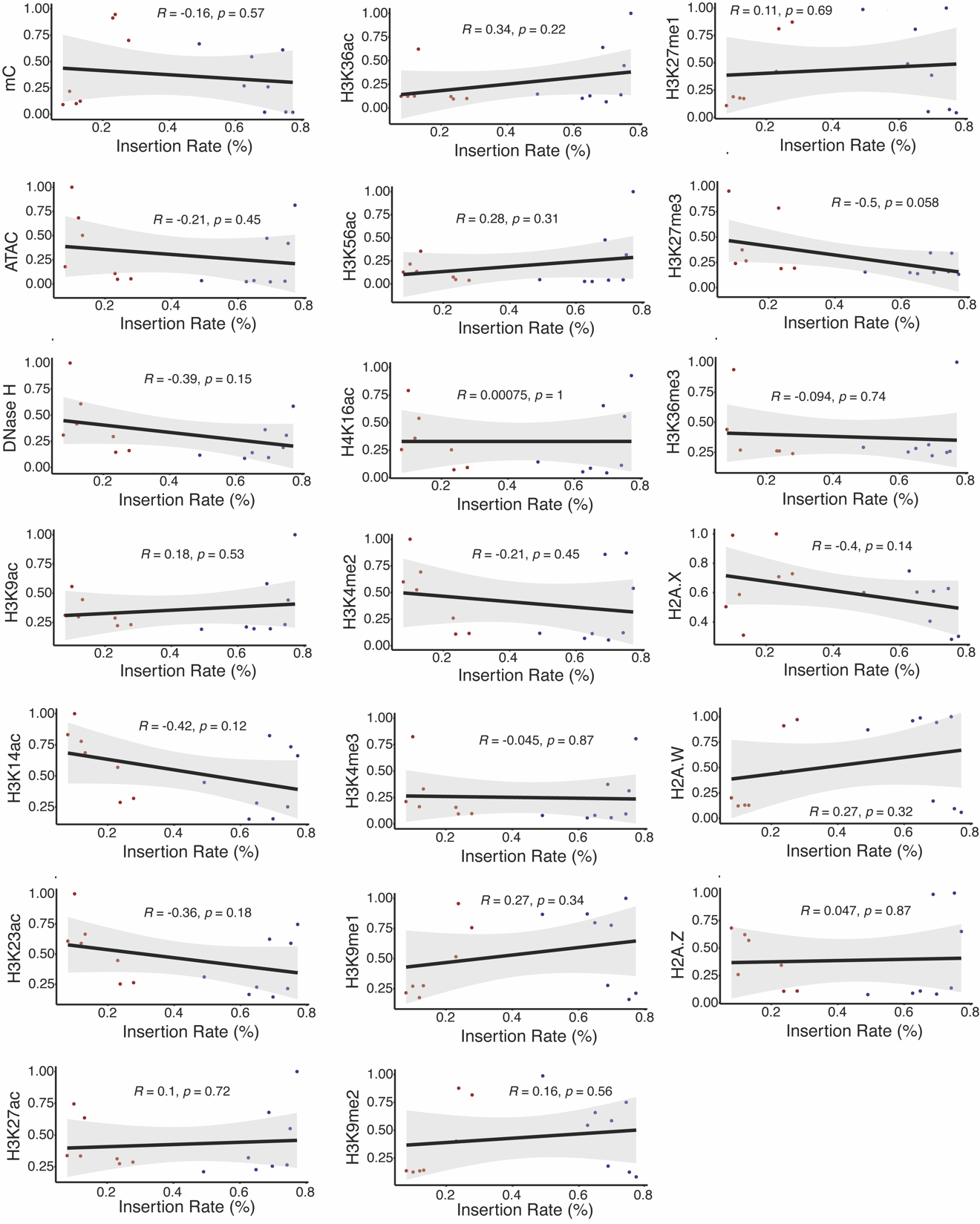

**Fig. S9.** Correlation analysis for 1 bp insertion rate and chromatin features. The blue dots represent MCsite4, and the red dots represent MCsite5. The trendline is black with gray indicating the standard error. The R value and *p*-values are indicated at the top of each correlation plot according to Spearman’s rank correlation coefficient. Each chromatin feature was normalized using all 15 target sites to allow for comparison between the CRISPR target sites on a scale of 0 to 1, with higher values indicating the higher levels for the respective chromatin feature.

Table S1. Primer sequences to amplify each CRISPR target site analyzed in these experiments.

**Table S2. Summary of NGS reads count for each tested target site.**

Read counts used to characterize mutagenesis frequency and mutation outcomes were shown in the “edited reads count” column for each site. The numbers were derived from the sum of all three replicates.

Dataset S1. Characterization of the sequences, DNA methylation, chromatin accessibility, and chromatin states for the 7,971 candidate CRISPR target sites identified.

Dataset S2. Characterization of the 7 candidate MCsite chromosomal locations, DNA methylation domain, chromatin accessibility, chromatin state, gene annotation, and RNA detection.

**Dataset S3.** Characterization of the 1 kb region (500 bp upstream and 500 bp downstream) flanking the CRISPR-Cas9 cut site. For each dataset, the values for each individual nucleotide flanking the CRISPR cut site in the 1kb window (500 bp upstream and 500 bp downstream) were quantified by calculating the sum.

**Dataset S4.** Index for matching samples with the correct fastq files.
